# Easing OMERO adoption with ezomero

**DOI:** 10.1101/2023.06.29.546930

**Authors:** Erick Martins Ratamero, Kiya Govek, Julio Mateos Langerak, Fernando Cervantes Sanchez, David J. Mellert

## Abstract

Many research laboratories need to manage, process, and analyze the increasingly large volumes and complexity of data being produced by state-of-the-art bioimaging platforms. OMERO is a popular open-source client-server application that provides a unified interface for managing and working with bioimages and their associated measurements and metadata. Integrating OMERO into analysis pipelines, such as those developed around the scientific Python ecosystem, will thus be a common pattern across the field of bioimaging. While OMERO has a powerful Python API, it provides minimal abstraction from the underlying OMERO object model and associated methods, which represent more complexity than most users are interested in for the context of an analysis script. We introduce ezomero, which was designed to provide a convenience layer on top of existing OMERO APIs and return data types that are either Python primitive or commonly used in scientific Python. Ezomero has minimal dependencies in addition to the OMERO Python library itself and is installable directly from PyPI. Here, we provide an overview of ezomero as well as several vignettes to illustrate how it can be used to accelerate discovery.

## Introduction

Increased adoption of large-scale imaging technologies, such as high content screening platforms, slide scanners, and light-sheet microscopes, has created a significant data management burden for many biological research laboratories and core facilities. The Open Microscopy Environment (OME; Goldberg et al., 2005) group developed OMERO (OME-Remote Objects; Allan et al., 2012) to provide a free and open source solution for this emerging challenge. In addition to being broadly adopted across the bioimaging community, OMERO has a stable and committed core development team, is interoperable with many other commonly used free and open-source software (FOSS) bioimaging tools (e.g., ImageJ, Rueden et al., 2017), and includes a rich default feature set as well as popular plug-ins that extend its functionality, e.g., OMERO.figure (The Open Microscopy Environment, 2023a) and OMERO.iviewer (The Open Microscopy Environment, 2023b). OMERO is also able to serve as the foundation for image data repositories such as the Image Data Resource (https://idr.openmicroscopy.org, Williams et al., 2017), a utility that will become more important as research funding agencies require all primary research data to be either deposited into public repositories or otherwise broadly shared with the research community.

One key advantage of OMERO is its well documented application programming interface (API) services, which enable developers and power users to both write clients that extend OMERO and script routine functions and workflows. Moreoever, although OMERO is a Java application, its API services can be invoked remotely from several other programming languages using bindings that are dynamically generated by Ice, an open-source remote procedure call framework developed by ZeroC. The OME group has also written a more user friendly Python module called BlitzGateway, which simplifies many common OMERO operations and wraps OMERO objects with a more “Pythonic” interface. Python developers are able to access OMERO’s Python bindings, BlitzGateway, and a command-line interface written in Python by installing the omeropy package (The Open Microscopy Environment and Glencoe Software, Inc., 2023), maintained by OME and available on the Python Package Index (PyPI). While omeropy is a powerful library, one of its core tenets is maintaining cross-language support with other OMERO APIs—while this simplifies maintenance and documentation, for example, it means omeropy cannot be tailored to Python’s strengths by definition.

The Research Information Technology group at The Jackson Laboratory manages multiple OMERO deployments, for which we have written a collection of scripts for executing common administrative tasks. We have also given a number of power-user/developer–focused workshops in which we teach how to interact with OMERO in the context of a Python script. Our experience with these development and teaching activities led us to two observations: 1) Some common OMERO tasks can be quite verbose when written in Python, even when using BlitzGateway, and 2) users that are proficient in using Python for bioimage analysis but who are not formally trained in computer science often struggle to work with the OMERO objects returned by omeropy. In addressing both of these issues, we developed over time a set of convenience functions that we felt provided a more comfortable Python interface. We ultimately bundled these functions into a Python package we call ezomero (The Jackson Laboratory, 2023).

Although ezomero initially grew organically with need, we made two early choices that allowed us to later formulate a coherent API. First, we decided to keep the use of OMERO- or ezomero-specific objects to a minimum, focusing instead on creating functions for which inputs and outputs are limited to either Python primitive types (e.g., strings, integers, etc) or objects familiar to users of NumPy or other scientific Python libraries (e.g., `numpy.array`). Second, we used function names that clearly explain the function’s purpose through the use of HTTP vocabulary (e.g., get, post, put) where applicable, and through consistent vocabulary (e.g., link to, print) where no obvious HTTP verb made sense. Here we provide an overview of the current state of ezomero, further develop our rationale for its use, and demonstrate how it can be used to accelerate the development of OMERO-based image analysis.

## Methods

### Source Code and Library Organization

Ezomero source code is available on Github (https://github.com/TheJacksonLaboratory/ezomero). We follow a simple Python package organization in which most functions are imported directly from the package root. Two modules, `ezomero.json_api` and `ezomero.rois`, contain functions for working with OMERO’s json API and regions-of-interest (ROIs), respectively. Despite the relatively flat package structure, the code has been split over multiple files (e.g., `_gets.py`, `_posts.py`,`_importer.py`) to improve readability and maintenance. In addition to the Python package directory, the code repository also contains separate directories for documentation (docs) and testing (tests). Ezomero source code is covered by the GPL-2.0 license, following the licensing of omeropy.

### Documentation

Ezomero is autodocumented by Sphinx (https://www.sphinx-doc.org), which builds html documentation from Python docstrings. We chose to follow NumPy docstring conventions, allowing us to autogenerate documentation using Sphinx’s napoleon extension (`sphinx.ext.napoleon`). We have published html documentation at https://thejacksonlaboratory.github.io/ezomero. All files necessary for Sphinx generation of documentation is located in the docs directory repository.

### Development Environment

As noted above, git/Github are used to manage ezomero version control. Users are invited to submit bug reports and feature requests using Github issues, and external contributions are welcomed. We have included a contribution guide as part of our repository, with guidelines for how to start contributing to ezomero, design principles that encapsulate the general direction we expect the project to pursue, and stylistic guidelines for consistency with existing code. A Code of Conduct is also provided, to detail the environment we would like to foster for collaboration and clearly delineate how interactions with users and contributors to our public Github repository should happen.

We use pytest for testing. Unit tests, pytest configuration, and some test data are found in the tests directory of the repository. When developing locally, we use docker-compose to deploy an ephemeral OMERO instance, which includes OMERO.server, OMERO.web, and a PostgreSQL database. However, we also run tests on pull requests and associated commits using Github actions, so contributors are not required to set up their own testing environment. We have configured some basic OMERO objects and test data as fixtures, but for the most part, objects are created as needed for a particular unit test on the ephemeral OMERO server and torn down when the test is complete.

### Installation and Deployment

The main dependency for ezomero is omero-py, the Python library for interacting with an OMERO server provided by OME. Ezomero also requires numpy for interacting with numerical arrays, and has pandas as an optional dependency for interacting with tables. Ezomero is available from PyPI, meaning a simple `pip install ezomero` is all that is required for installation, provided dependencies are available. Trying to automatically install ezomero from PyPI without the requisite dependencies can be problematic; in particular, omero-py depends on ZeroC-Ice’s python library, which is not readily available from PyPI as a prebuilt package; Ice is built statically at install time. This process is time-consuming and a source of errors. As an alternative, prebuilt ZeroC-Ice packages can be installed via conda (from the condaforge channel), and ezomero installation can be completed with `pip`. Full installation instructions are available on the ezomero GitHub repository.

## Results

### Design

As noted in the introduction, ezomero was designed to minimize user interaction with OMERO model objects and to make routine administrative tasks simple to code. In this way, ezomero is meant to complement rather than replace the APIs available through omero-py. If a skilled developer intends to write highly efficient code that makes sophisticated use of the OMERO model objects, ezomero may not be a good solution—omero-py offers, for example, direct database querying options that can provide more complex functionality that ezomero does not aim to replicate. Our typical user is, for example, a graduate student that wants to analyze their image data stored in OMERO with scikit-image (Van der Walt et al., 2014) or torchvision (Marcel and Rodriguez, 2010) using a Jupyter Notebook. Another potential user might be a core facility director that needs to write a script to perform a nightly administrative task. These users may want to write code quickly, without relying too heavily on OMERO documentation.

Use of ezomero begins with the creation of a session for connecting to OMERO.server. A BlitzGateway object (provided by omero-py) is used for the connection; we typically name this object `conn`, which is consistent with OME’s own examples. While users can create the BlitzGateway object directly, we have provided a convenience function, `ezomero.connect`, that can simplify the creation of this object by reading from stored connection parameters or environment variables, or prompting the user to input parameters, including a password, that were not passed to the function. Nearly all other ezomero functions take the `conn` object as the first parameter.

We have used HTTP vocabulary in our function names in an effort to make code more readable, with obvious purpose. We use “get” whenever data needs to be retrieved from the server and “post” whenever a function creates a new resource on the server. We use “put” in one case, `ezomero.put_map_annotation`, which will replace the key-value pairs in an OMERO map annotation. Other common verbs are “print” for displaying various kinds of information to Stdout, “filter” for narrowing lists of OMERO Image IDs, and “link” for creating Project, Dataset, and Screen links that are used to organize Images.

### Example use cases

This section aims to show the main advantages ezomero presents over the Blitz API for novice programmers and image analysts: a simplified, less verbose interface with an OMERO server, and usage of standard scientific Python data structures as inputs and outputs.

To do so, we will present a few 1-to-1 comparisons for simple tasks and discuss how the code for ezomero differs from Blitz API code.

The first 1-to-1 comparison we want to present is a very simple workflow that might precede data analysis, consisting of creating a connection to an OMERO server and retrieving a 5-dimensional image as a numpy array. The ezomero workflow tracks our natural language description of the problem pretty closely. Listing 1 shows us that: the workflow is simply 1) connecting to the server, and 2) retrieving an image. These actions are a single line each. Additionally, the connection parameters are either retrieved automatically by ezomero, and/or requested from the user at runtime.

On the other hand, the same pipeline using the Blitz API, presented on Listing 2, requires 1) creating a connection object from a CLI client, 2) retrieving a Blitz gateway–wrapped `Image` object from the server, 3) getting the wrapped `Pixels` object, 4) getting the dimensions of the image, 5) creating a list of planes, and 6) iterating over that and passing it to `getPlane` to achieve the same result.

**Listing 1.**
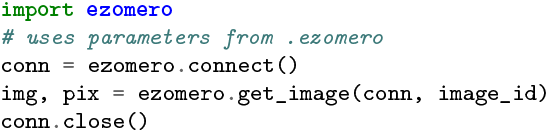
Connecting to an OMERO server and retrieving a 5d image to a numpy array using ezomero.

**Listing 2.**
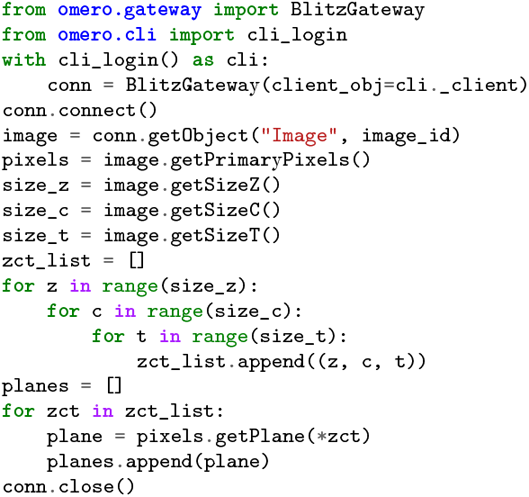
Connecting to an OMERO server and retrieving a 5d image to a numpy array using the Blitz API.

Our second featured use case is a simple administrative workflow, where a user (or an administrator) wants to create a new Dataset inside an existing Project in an OMERO server.

Again, the ezomero version presented in Listing 3 tracks our natural understanding of the problem, requiring 1) a single line for connecting to the server, and 2) a single line for posting a Dataset (with an optional project_id parameter).

The same workflow using the Blitz API version is shown in Listing 4. We see that this verbose process requires: 1) a new `DatasetI` object to be created, 2) the Dataset’s name set, 3) the connection object’s `UpdateService` retrieved and used to save the `DatasetI` object.

That only solves half the problem. We then need to: 4) create a `ProjectDatasetLinkI` object, and finally 5) set the Child and Parent attributes of the `ProjectDatasetLinkI` object before saving. Only then will the Dataset be correctly linked to the intended Project.

**Listing 3.**
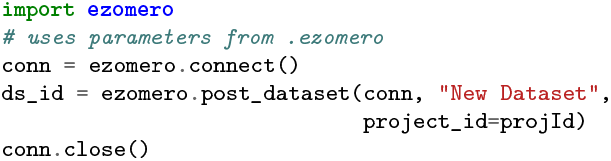
Connecting to an OMERO server and adding a new dataset to an existing project using ezomero.

Our third and last proposed use case extends our first example. Here we want to attach a dictionary as key-value pairs to a specific image from a Dataset with a key-value pair “analyze: true”. Such dictionary is generated by an `analyze_data` function applied to that specific image.

Once more, we find that the ezomero version presented in Listing 5 follows a simple, intuitive flow that closely tracks our description of the problem. Here, we 1) retrieve the image IDs on that dataset, 2) filter them by key-value pair to find the image of interest, 3) retrieve the image, 4) pass it to `analyze_data`, and 5) create a map annotation with the results.

**Listing 4.**
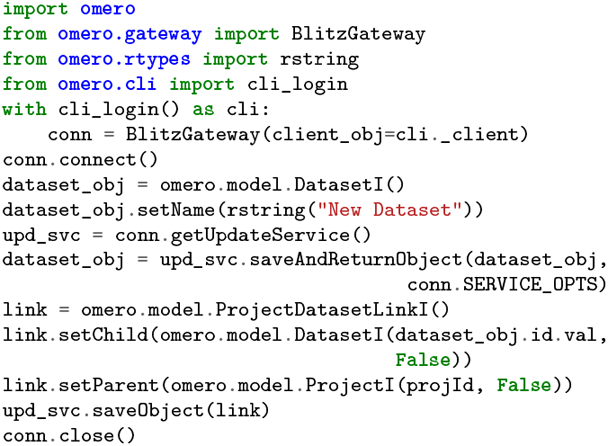
Connecting to an OMERO server and adding a new dataset to an existing project using the Blitz API.

The Blitz API version presented in Listing 6, again, is more complex and verbose. We need to 1) retrieve a `Dataset` object, 2) iterate over its children to get the image IDs linked to that dataset, 3) use the `QueryService` from the connection object to create an HQL query that returns the image ID with the desired key-value pair, 4) use a structure similar to Listing 2 to retrieve the full image and pass it to `analyze_data`, 5) reshape the returned Python dictionary into a list, 6) create a wrapped `MapAnnotation`, 7) set the namespace and value of the `MapAnnotation` object, and 8) save the `MapAnnotation` and link it to the correct image.

**Listing 5.**
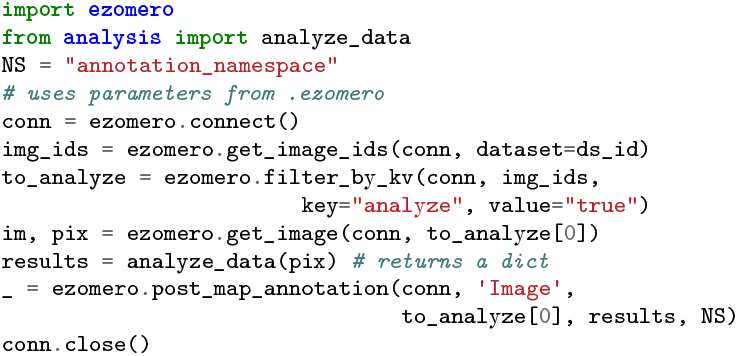
Using ezomero in an analysis workflow.

These use cases cover a range of relatively simple workflows for users and administrators interacting with an OMERO server, and in each one of them we can see how ezomero leads to fewer lines of code and lesser cognitive load. It allows for cleaner, more easily maintainable code, and decreases the necessary knowledge of the full OME data model and OMERO Blitz API significantly.

**Listing 6.**
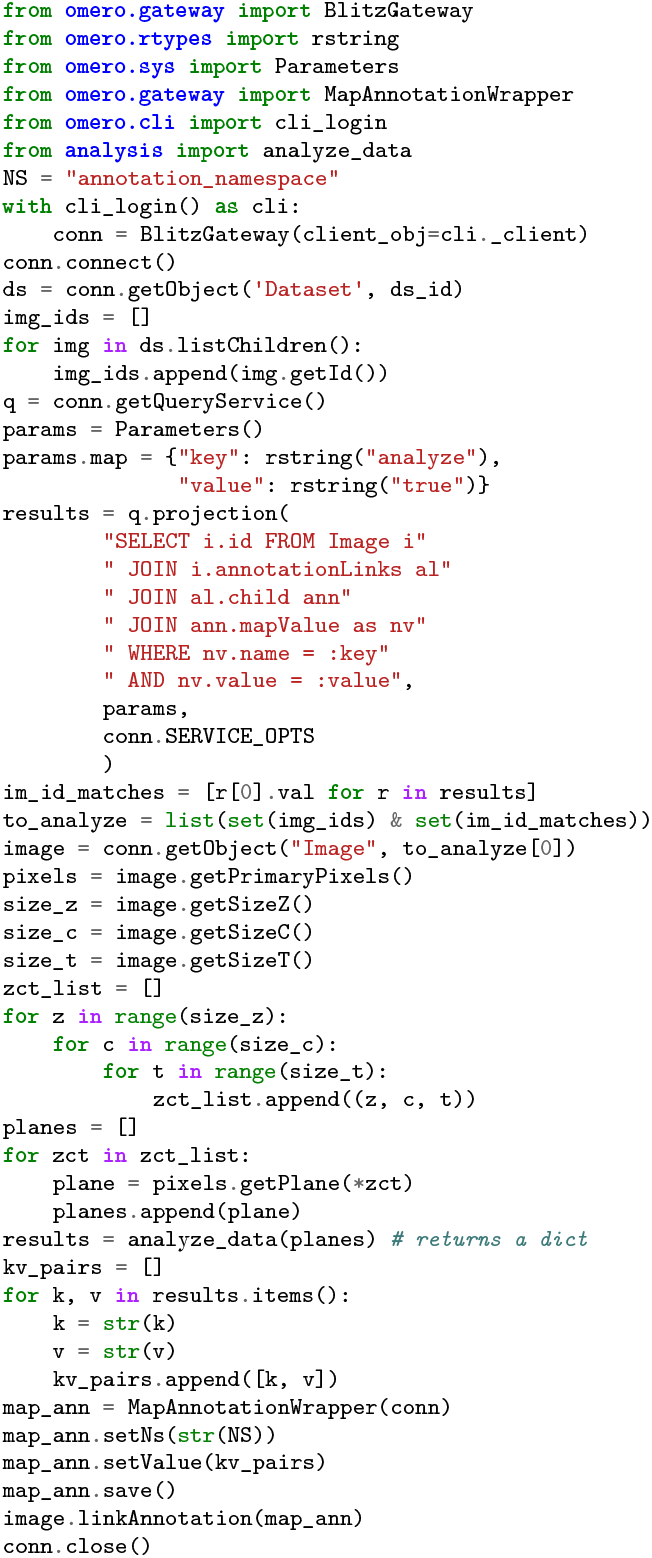
Using Blitz API in an analysis workflow.

## Discussion

We described ezomero, a convenience-layer library for interacting with an OMERO server using Python. This library, by simplifying code and allowing for a lower barrier of entry for end-users and administrators interested in programmatic access to the OMERO platform, can help ease OMERO adoption in institutions and make more sophisticated workflows possible.

Our initial impetus for creating the functions that eventually developed into ezomero was a desire for automation and reproducibility in our day-to-day OMERO tasks. We believe this to fill an important niche in the imaging data management community, decreasing the necessary technical expertise for creating code-based workflows that can be shared, reviewed and developed upon more easily.

Importantly, OMERO itself is not static: it is also software that is under constantly development. We believe the OMERO platform is uniquely positioned to be a future-proof data management platform, becoming more a metadata repository and discovery layer, and aggregating distributed data across on-premises and cloud storage sources. In this environment, programmatic access will be more important than ever, and creating tools to facilitate such access is paramount.

## ACKNOWLEDGEMENTS

The authors gratefully acknowledge the OME team for the years of fruitful collaboration and thoughtful discussion, in particular Josh Moore and Will Moore for the thoughtful feedback on this manuscript.

## AUTHOR CONTRIBUTIONS

Conceptualization, E.M.R. and D.J.M.; Methodology, E.M.R. and D.J.M.; Software, E.M.R., K.G., J.M.L., F.C.S. and D.J.M.; Investigation, E.M.R. and D.J.M.; Writing and Original Draft, E.M.R. and D.J.M.; Review and Editing, E.M.R., K.G., J.M.L., F.C.S. and D.J.M.; Funding Acquisition, E.M.R. and D.J.M.; Resources, E.M.R. and D.J.M.; Supervision, E.M.R. and D.J.M.

